# Network analysis of high-density microelectrode recordings

**DOI:** 10.1101/139436

**Authors:** Torsten Bullmann, Milos Radivojevic, Stefan Huber, Kosmas Deligkaris, Andreas Hierlemann, Urs Frey

## Abstract

A high-density microelectrode arrays (HDMEA) with 3,150 electrodes per square millimetre was used to capture neuronal activity across various scales, including axons, dendrites, and networks. We present a new method for high-throughput segmentation of axons based on the spatial smoothness of signal delays. Comparison with both ground truth and receiver operator characteristics shows that the new segmentation method outperforms previous methods based on the signal-amplitude-to-noise ratio. Structural and functional neuronal network connectivity were reconstructed using a common extension of “Peter’s rule” and a inter-spike histogram method, respectively. Approximately one third of these connections are putative chemical synapses. We evaluated the spike patterns but did not find evidence for “polychronisation” (non-synchronous but precisely timed spike sequences). The developed framework can be used to investigate the relationship between the topology of neuronal connections and emerging temporal spike patterns observed in dissociated neuronal cultures.

## 1 Introduction

A key problem in neuroscience is to understand the relationship between structure and function. The combination of diffusion magnetic resonance imaging and functional magnetic resonance imaging can be used to address this problem at the macroscopic level [57][25]. Recent efforts, such as the BRAIN Initiative (US), the Human brain project (EU) as well as the Brain/MINDs initiative (Japan) aim at the development of new technologies for mapping the functional connectome and large-scale recording of neuronal activity. But even for more accessible model systems, such as cultured dissociated primary neurons the relationship between the topological structure of the neuronal networks and the emerging temporal activity patterns is not easy to understand. One reason is that the structural connectivity graph, which is given in neuronal network simulations, is usually not known for experimental data of network activity.

We propose that high-density microelectrode arrays (HDMEAs) could be used for the high-throughput acquisition of single neuron electrical footprints revealing their axons and dendrites according to their typical waveform [47] and this information could be used for the subsequent reconstruction of their connectivity according to an extension of “Peter’s rule” [46][8]: the probability of two neurons being connected is proportional to the overlap of their axonal and dendritic fields [50].

To evaluate the applicability of this concept, both sub-cellular and network-wide activity recordings were acquired at the same time from rat cortical neurons cultured at very low density on HDMEAs. After identification of the axon initial segments (AISs), action potentials were tracked across full axonal arbours of single neurons at sub-cellular resolution, and the dendritic field of single neurons was inferred by the positive return current during action potential generation. The data showed that structural connectivity of a small network overlaps with the functional connectivity obtained from pairwise inter-spike interval histograms.

In the present study, HDMEAs and optical imaging of primary rat cortical cultures were combined in order to establish a framework for analysing the relationship between the topological structure of the neuronal networks and the emerging temporal activity patterns, observed in dissociated neuronal cultures, including: (1) considerations for efficient acquisition of network activity at sub-cellular resolution, (2) segmentation of axons by considering the neighbourhood standard deviation of the transmission delay, (3) segmentation of dendrites according to their positive signal peak, (4) discrimination of GABA vs. glutamatergic neurons by peak width, (5) structural connectivity inference by determining the overlap between axonal and dendritic fields, (6) inference of functional connectivity according to inter-spike intervals, (7) network characterisation by graph invariants, (8) comparison between structural and functional network connectivity, (9) detection of putative chemical synapses, and (10) detection of spike patterns.

The HDMEA used in this study featured 11,011 electrodes at a resolution of 3,150 electrodes/mm^2^ (pitch of 17.8 *µ*m) [18]. It employs the switch-matrix concept [45], to provide high signal quality at very low noise levels. However, the number of parallel recording channels was limited to 126 channels connected to flexibly selectable subsets of electrodes in so-called “configurations”.

## 2 Results

To initially identify the location of AISs, the whole array was scanned by using configurations in which non-overlapping blocks of 6×17 electrodes were connected to the amplifiers through the switch matrix [18]. A local maximum in the amplitude of the negative voltage peak correspond to putative (proximal) AIS locations [5]. In the low-density cultures up to 100 AISs could be distinguished. So called “fixed electrodes” were selected as trigger electrodes at the putative locations of the AISs, while the remaining “variable electrodes” were selected in sequential configurations (see next section) to map the axonal arbours over the entire array. For the selection of fixed electrodes all electrodes were ranked according to their median negative peak amplitude. The electrode with the highest rank was selected, afterwards all electrodes in its proximity (within 60 *µ*m distance) were discarded from the list, and the procedure was repeated. Taking advantage of the large negative peak amplitude near the AIS, the threshold for event detection was set to 6 times the noise level. This prevented crosstalk from other neurons while still reliably detecting the activity of that single neuron.

### 2.1 Optimal recording configurations for high-throughput scanning

The extracellular signals originating from axons and dendrites are very small with respect to the background electrical activity and noise, so that spike-triggered averaging must be applied. We developed a set of recording configurations to map the electrical footprint of several neurons in parallel by utilising the switch matrix of our HD-MEA. The switch matrix can be dynamically configured to connect a large number *e* of electrodes to a smaller number *a* of amplifiers. A naive method would use one electrode as trigger and the remaining electrodes to scan the neuronal footprint, which results in a large number of configurations *c_w_* needed to scan the whole array, *c_w_* = *e/a*. If we instead intend to record axonal arbours of *n* neurons, electrodes near the AISs of these neurons have to be always connected, each to one amplifier (*n* fixed electrodes). The remaining amplifiers can then be connected to the remaining electrodes in successive configurations (variable electrodes). In this way the whole array can be scanned in

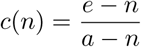

configurations. For *n* neurons we need *c*(*n*) configuration, which means on average *C* (*n*) = *c*(*n*)*/n* configurations for one neuron. An optimal strategy means to choose *n* such that *C* (*n*) → min for 0 < *n* < *a*. With *n* < *a* ≪ *e*, the number of neurons being much smaller than the number of electrodes, we can approximate by:

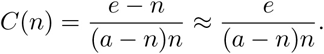

The right hand side has a minimum for *n* = *a*/2. Therefore, approximately half of the amplifiers should be connected to fixed electrodes. The other half of the amplifiers can then be used to scan the whole array in *c* ≈ *e*/(*a*/2) = 2*c_w_* configurations, or on average *C* ≈ *e*/(*a*/2)^2^ = 4*e/a*^2^ configurations per neuron. The axonal arbours of a single neuron may extend over the whole array, but by scanning the axonal arbours of many neurons in parallel, the average number of configurations per neuron is much less than required for scanning the whole array *C* ≈ 4*e/a*^2^ ≪ *c_w_* = *e/a* for large *a*. This is because increasing the number of amplifiers decreases quadratically the average time to scan a single neuron, which allows for high-throughput acquisition of axonal delay maps.

### 2.2 Identification of axonal arbours

Previously (see Figure 6 in [4]), axonal arbours were traced according to the negative peaks in the spike triggered averages that exceeded 5 times the background noise (*s_V_*). This method (hereafter termed “method I”) evaluates the spike-triggered averages for each electrode separately, without considering their spacial arrangement and correlation between neighbouring electrodes. Interestingly, more realistic axonal contour can be drawn by humans in visual observations of the spatial movement of signal peaks in movies [49]. This motivated us to explore several methods of computer vision to segment axons after extraction of the axonal signals, e.g. Markov random fields and optical flow, and compared their results with a ground truth as well as with method I. It turned out, that a simple method (hereafter “method II”), based on thresholding the “spatial smoothness” of the delay of the negative peak is fast and reliable. When we mapped the delay of the negative peaks present in the spike triggered averages this map showed a distinct region of locally “smooth” delays against a background of random delays (Figure 1b). This is due to the fact that signals originating from a common source, e.g. from axons of the same neuron, their delays are very similar at neighbouring electrodes, whereas for a random signal the negative peak could occur anywhere in the interval [Δ*t_pre_*, Δ*t_post_*] for which the spike triggering was performed. Δ*t_pre_* and Δ*t_post_* represent the boundaries relative to the spike trigger, thus the total spike triggered average has the length *T* = Δ*t_post_* − Δ*t_pre_*. To quantify the smoothness of the delay map the delay was sampled in the Neumann neighbourhood (compare Figure 2h) around each electrode and the sample standard deviation 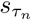 was calculated. Note, that in the case of a uniform distribution over the interval [0, 1], the standard deviation of that sample is bounded by 0 ≤ *s* ≤ 0.5 and shows a characteristic distribution depending on the number of observations *N* in each sample. This distribution shows a sharp peak around at the mean standard deviation:

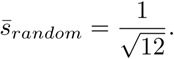

There is no analytical expression for this distribution, but after a coordinate transformation of the interval it can be approximated by a beta distribution 𝓑(*α, β*).

**Figure 1:**
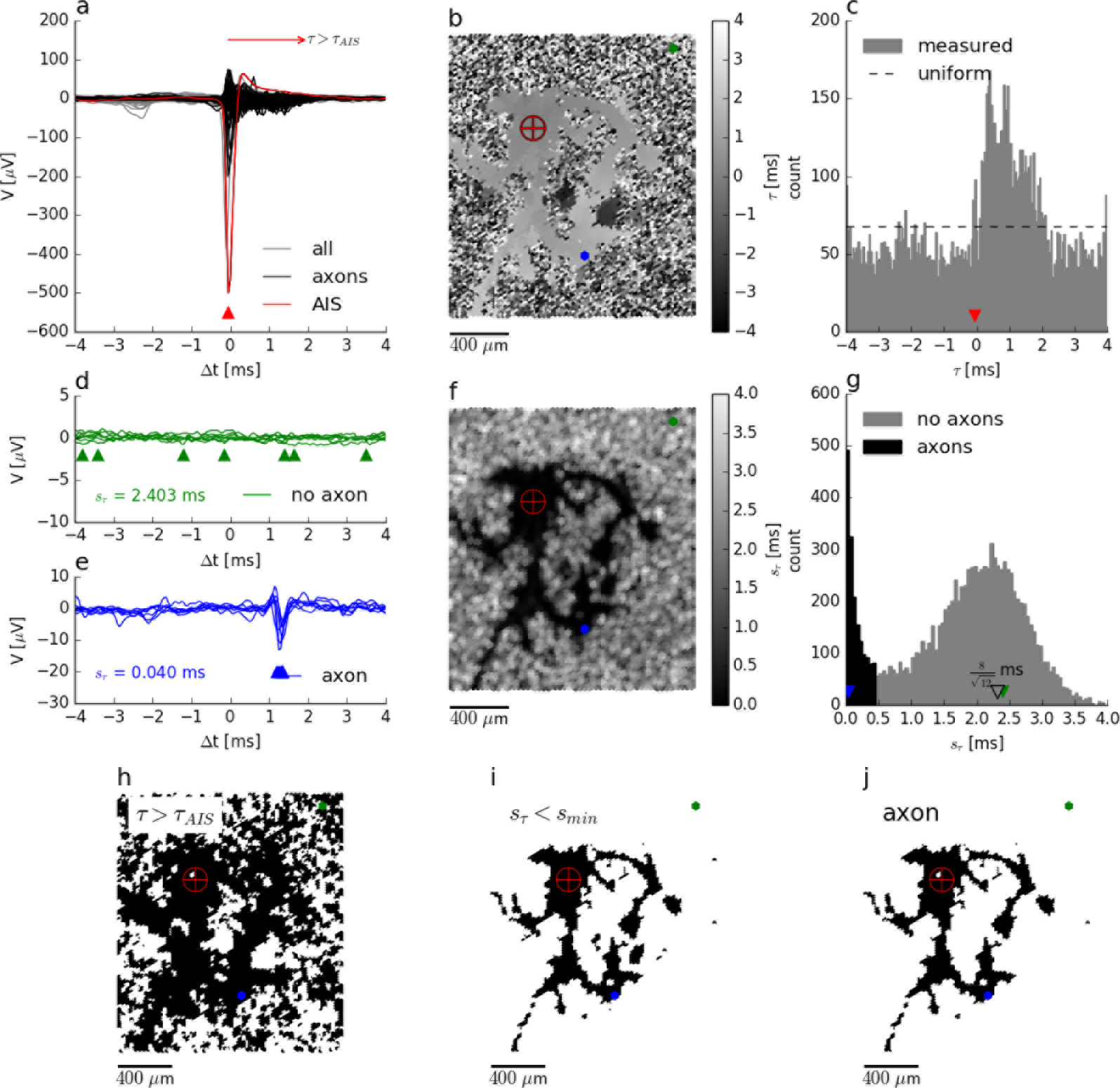
Segmentation of the axonal arbour based on the spatially correlated spontaneous activity of a single neuron. Spike-triggered averages **(a)** for electrodes located close to the (proximal) axon initial segment (AIS) (red trace), close to axons (black) and recording background activity and noise (grey). The negative peak at the AIS appears slightly earlier than at the trigger electrode. Mapping **(b)** and histogram **(c)** of the delay of the negative peak, *τ*, showing an irregular shaped area with a “smooth” grey value outlining the axonal arbour, which is surrounded by a “salt-and-pepper” patterned background area. Axonal signals appear at 0 ms < *τ* < 2 ms. Spike-triggered averages for *N* = 7 neighbouring electrodes located in the “salt-and-pepper” region **(d)** have a large sample standard deviation for the delays, 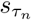, as compared to those located in the “smooth” region **(e)**. Mapping **(f)** and histogram **(g)** of 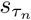. The small irregular shaped area outlining the axonal arbour is dark, whereas the surrounding area features lighter colors. Segmentation is done by placing the threshold *s_thr_* ≈ 0.5 ms in the valley between the sharper peak (black) close to 0 ms and the broad peak (gray) around the expected 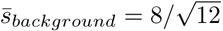 ms (open triangle) for random delays. Mapping of electrodes where the negative peak appears after the negative peak of the AIS **(h)**, with 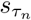 < *s_min_* **(i)**, and presumably axonal signals**(j)**, corresponding to the intersection of both. **Legend:** The crosshair symbol shows the location of the (proximal) AIS, the green and blue dots represent a patch of 7 neighbouring electrodes located in the “salt-and-pepper” and “smooth” areas, respectively. Corresponding negative peaks are indicated by triangles of the same colour.

In case where axonal signals are present, for each electrode with a negative peak at *t* the delays of its neighbourhood are distributed in the interval [*t* − *r/c*; *t* + *r/c*] depending on the velocity *c* of the action potential and the distance *r* between the electrodes. Therefore, we can assume a similar distribution of the sample standard deviation for axonal delays distributed around the mean standard deviation:

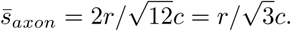

For a HDMEA with *r* = 18 *µ*m and a typical conduction velocity for short range projecting axons in rat neocortex of 0.3 − 0.44 m/s [37][62] a standard deviation around 30 *µ*s can be expected. At the boundary, more and more neighbouring electrodes do not pick up the axonal signal and the sample standard deviation shifts towards

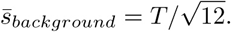

Empirically, the distribution of *s_axon_* can be approximated by an (truncated) exponential distribution *ε*(*γ*) (see below). Therefore, axons have distribution of 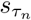 with a peak close to zero and which is clearly distinguishable from the distribution for the background. A threshold *smin* placed at the local minimum in the 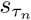 distribution (Figure 1g) separates both populations. The electrodes with a 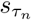 below this threshold represent negative peaks that are consistent across neighbouring electrodes (Figure 1i). If these peaks appear after the negative peak at the AIS (Figure 1h) they are assumed to originate from the axonal arbour of the neuron (Figure 1j).

For a limited number of neurons the ground truth was available and we could compare the new method (method II) with the method I employing a fixed threshold at 5*s_V_* [4]. An example of an transfected neuron is shown in Figure 2c. More electrodes were selected by method II than by method I, which could be compensated by lowering the threshold, e.g. to 3*s_V_*. In order to compare the electrode selection *E* with the ground truth axon label *A*, we used the Hausdorff distance *H*, which is commonly used in computer vision to measure how far two shapes are from each other. Thus, after registering the label image to the electrode coordinate system, we calculated:

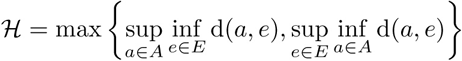

using the Euclidean distance d(*a, e*) between the electrode *e* recording an axonal signal and the pixel *a* labelled axon. Method I showed a smaller deviation from the ground truth than method II. This was mainly due to the fact that by decreasing the threshold, more electrodes far away from the axonal arbours were selected. The new method, inherently relying on the adjacency, rejected these “outliers” and produced more compact maps that more closely followed the ground truth. In other words, the distributions for activity classified as axonal signals and as background show a larger overlap for method I than method II. We tested the robustness of method II against the spatial distance of the electrodes by increasing the spatial extension of the neighbourhood while keeping the number of electrodes in each neighbourhood constant (N=7). When hexagonal patterns with *r* = 2 × 18 *µ*m or *r* = 3 × 18 *µ*m distance between electrodes was selected, the distance to the ground truth only slightly increased (Figure 2g). However, it seems that for a carefully chosen threshold (e.g. around 4.5*s_V_*, 𝓗 ≈ 100 *µ*m, Figure 2f), the results of the original method perform as well as the new method.

**Figure 2:**
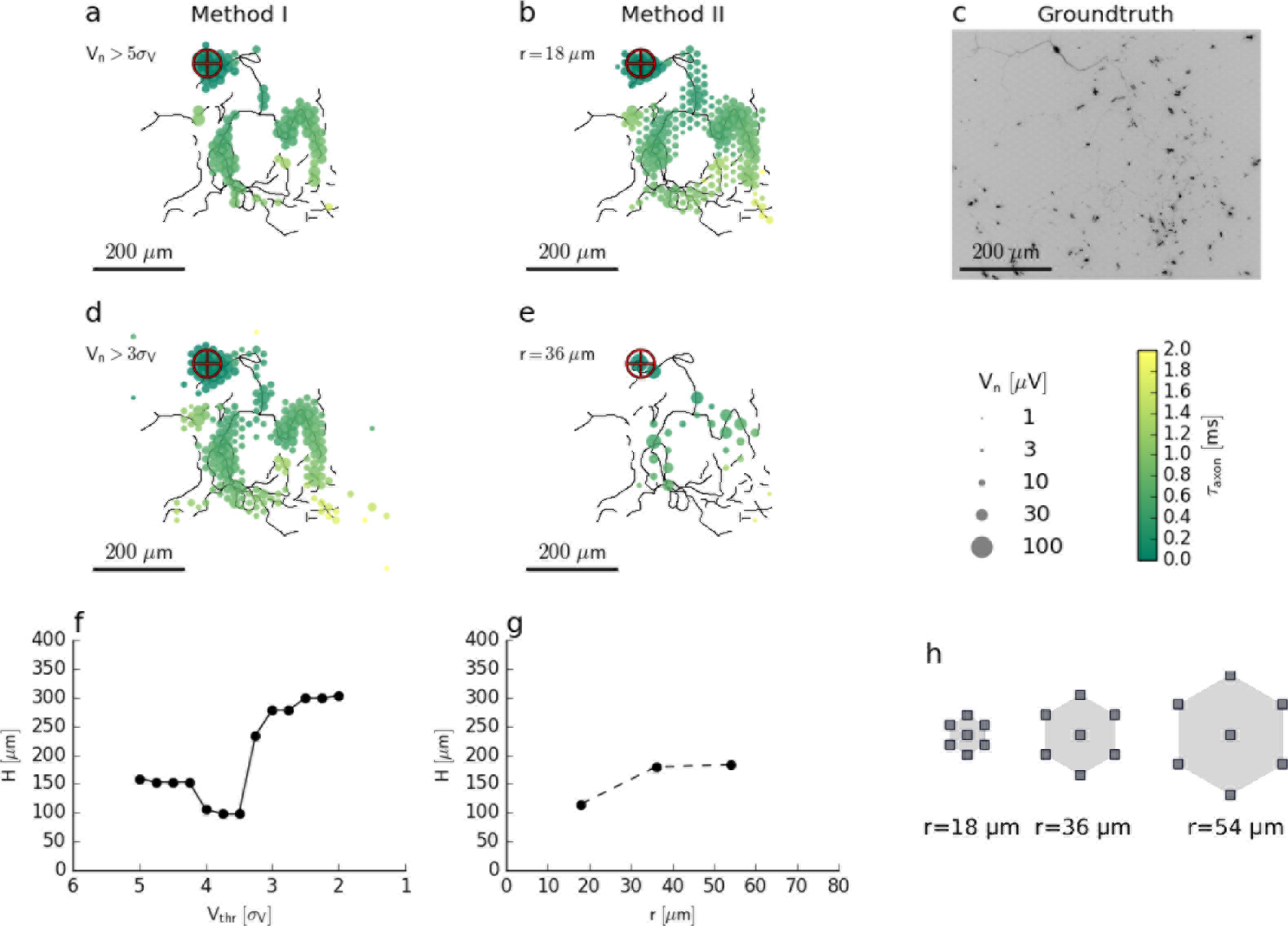
Evaluation of axon segmentation based on ground truth. Mappings and Hausdorff distance 𝓗 are shown for method I **(a, d, f)** and II **(b, e, g)**. The high threshold employed by method I leads to a higher false negative rate and a larger 𝓗 ≈ 150 *µ*m compared with 𝓗 ≈ 100 *µ*m for method II. Lowering the threshold from 5*s_V_* **(a)** to 3*s_V_* **(d)** for method I leads to more false positive electrodes far away from the axon (black outlines) and 𝓗 ≈ 300 *µ*m. In contrast, method II is robust **(g)** to an increased electrode distance **(h)**: increasing the distance from *r* ≈ 18 *µ*m **(b)** to *r* ≈ 36 *µ*m **(e)** ensures a higher true positive rate and 𝓗 ≈ 200 *µ*m. Axons were manually traced from fluorescence images (DsRed, LUT inverted) **(c)**.

Both segmentation methods are binary classifiers, and we compared them using their receiver operator characteristic (ROC)[17]. We fitted the respective empirical distributions of their scores with a mixture of two partially overlapping distributions representing axonal signals (“positive” class, P) and background activity (“negative” class, N):

1. two normal distributions (Figure 3a):

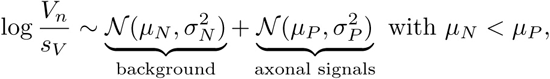
2. a beta distribution and a truncated exponential distribution (Figure 3b):

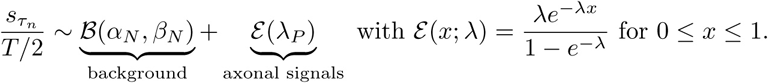

For each threshold, true positive rate (*TPR*) and false positive rate (*FPR*) were calculated from the confusion matrix and plotted as a ROC curve (Figure 3c). The area under the curve (*AUC*) showed that the new method consistently has a better performance than the original method for a total *n* = 23 neurons (Figure 3f). Furthermore, the automatic threshold procedure yields a much better *TPR* at the expense of a slightly increased false positive rate *FPR* compared with the original method (Figure 3g).

**Figure 3:**
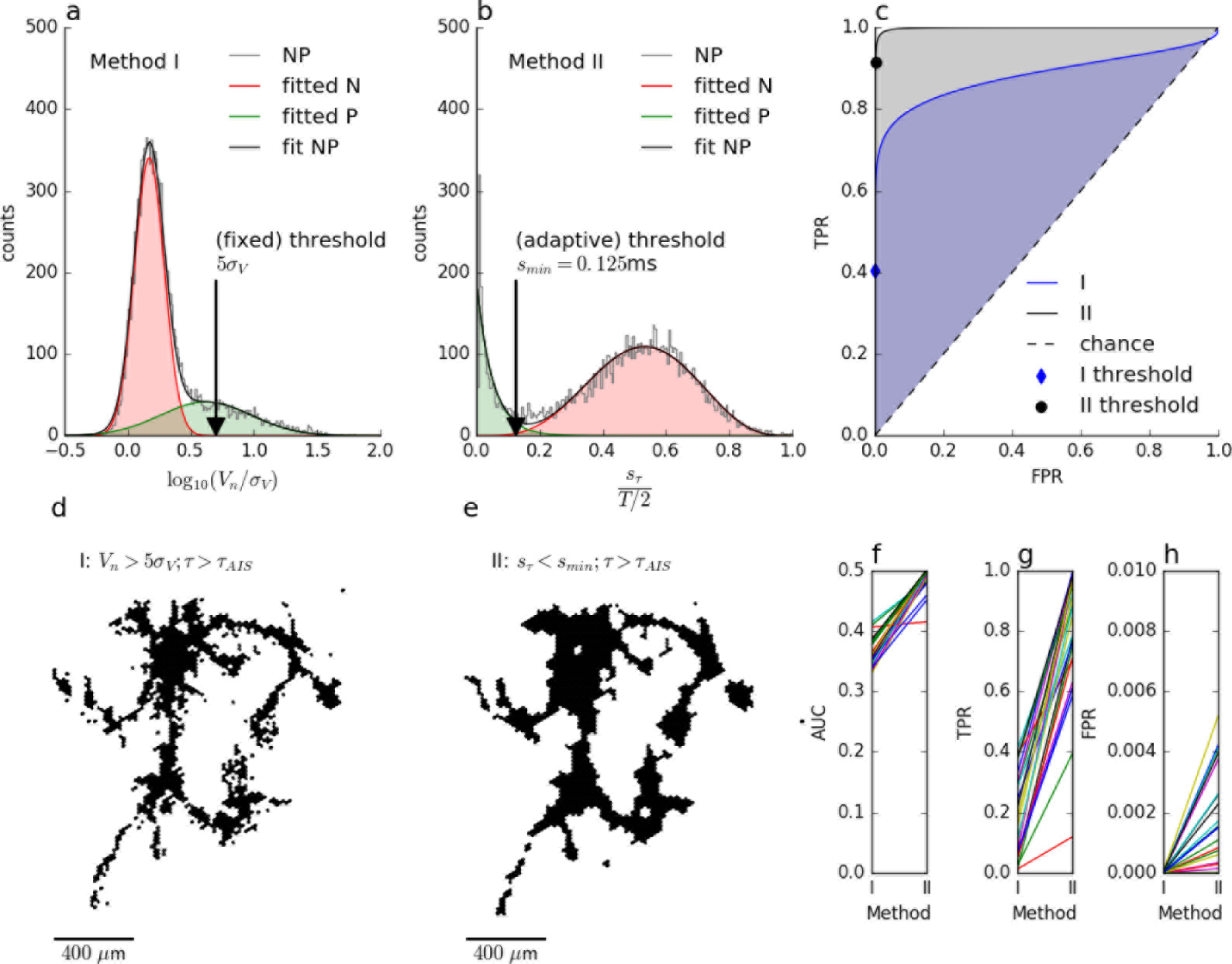
Comparison of the axon segmentation methods based on the receiver operator characteristic (ROC). Distributions and mappings are shown for the segmentation of an individual neuron segmented by method I **(a, d)** and II **(b, e)**. The empirical distribution (NP) of amplitude *V_n_* (log normalised by signal noise, *σ_V_*) and the sample standard deviation of the delay 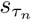 (normalised by *T*/2) of the negative peaks were fitted (fitNP) to obtain the distributions of axonal signals (positive class, P) and background activity (negative class, N). Respective true positive rate (TPR) and false positive rate (FPR) were calculated for each possible threshold and plotted as ROC curve **(c)**. The cross depicts the position of the (fixed) threshold of method I (FPR=0.00009, TPR=0.7), whereas the circle indicates the (adaptive) threshold of method II (FPR=0.011, TPR=0.85). Method II (blue shading) performs better than method I (grey shading) as shown by the larger area under the curve (AUC). This was true for all *n* = 23 neurons **(f)**, and, although method I has a lower FPR **(h)**, its TPR **(g)** is much lower than that of method II, as it misses more than 50% of axonal signals.

### 2.3 Dendritic arbours

In a rough approximation, the dendrite reveals itself by the positive extracellular potential [47] generated by passive and active properties of the dendrites. The passive component is associated with the sodium influx at the AIS, which leads to a return current, which charges the membranes and its amplitude is proportional to the membrane capacitance and inversely proportional to the electrotonic distance. To determine the time point of the sodium influx at the AIS, the full width at half maximum amplitude of the negative peak (|*δ_h_* |, commonly FWHM, full width at half peak) is used. It is given by the time interval the voltage trace crosses the half maximum amplitude right before and after the negative peak at the AIS:

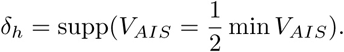

The map of the positive peaks that occur during the negative peak in the extracellular signal from the AIS (*V_AIS_*) shows the spatial extension of the dendritic field (Figure 4). Although it is clear that larger amplitudes correspond to dendrites whereas smaller amplitudes could also originate from axons (Figure 1e), it is difficult to determine an exact threshold that could be used to distinguish between them. Nevertheless, if the positive peaks appear during the half negative peak *δ_h_* at the AIS (Figure 1g) it can be assumed to originate from the dendrite(Figure 4f). Note that, although the estimation of connectivity depends on dendrites and axons, the spreading of the axon is the major defining factor, especially in case of long-range axons. Therefore, an estimation of the dendrites is less important.

**Figure 4:**
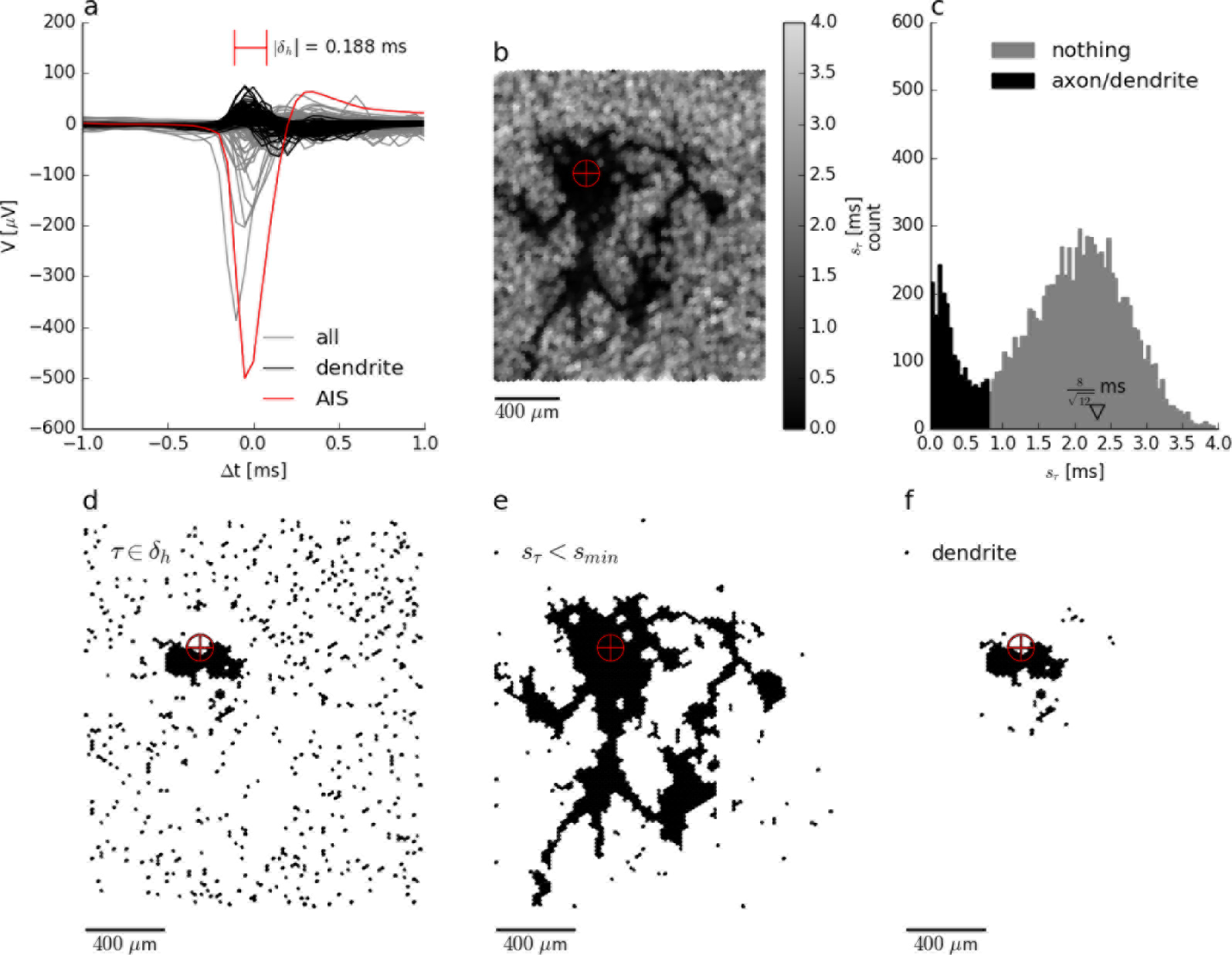
Segmentation of the dendritic arbour of a single neuron based on the return current generated during action potential initiation. Spike-triggered averages **(a)** for electrodes located close to the proximal fraction of the axon initial segment (AIS; red trace), close to the dendrite (black) and recording background activity (grey). The return current appears as positive peak during the negative peak at the AIS (red trace). Mapping **(b)** and histogram **(c)** of the sample standard deviation for neighbouring positive peaks, 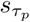. Note that axonal signals also possess a positive signal component and small peak, therefore smaller values of 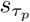 may represent to both, dendrites and axons (dark), whereas larger values around the expected value of 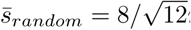 (open triangle) indicate random delays. Segmentation is performed using a threshold *s_thr_* ≈ 0.8*ms*, which is slightly higher than that used in (Figure 1G). Mapping of electrodes for which the positive peak appears during the half width of the negative peak at the AIS **(d)**, electrodes with with 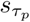 < *s_thr_* **(e)**, and electrode with presumptive dendritic signals**(f)**, corresponding to the intersection of both. **Legend:** The crosshair symbol shows the location of the (proximal) AIS.

### 2.4 Estimation of structural and functional connectivity

Structural connectivity is reconstructed according to a extension [50] of “Peter’s rule” [46][8], that the probability of two neurons being connected is proportional to the overlap of their axonal and dendritic fields (Figure 5a,b,c). It is possible to apply this rule on our data, because we segmented axonal and dendritic fields from the electrical footprint [47] of each individual neuron.

If the overlap (area) between the axonal and dendritic fields was larger than a threshold *ρ*, we assumed a “structural connection” (instead of “apposition”). The axonal delay *τ_axon_* was calculated as the median of the delays within the overlap.

There are several measures of functional connectivity, such as cross correlation, Granger causality, and transfer entropy, but these are derived for continuous time-series data (spike rate, calcium imaging) and lack robustness against bursting activity, which is present in cortical cultures (but see [59]). Therefore, we used a metric that is based on the relative timing of individual action potentials of two neurons (Figure 5d,e,f). Briefly, we calculated the probability distribution *P* for time lags *τ_lag_* between individual spikes of two neurons (inter-spike interval histogram, ISIH). Usually, the bursting dynamics results in a broad peak around *τ_lag_* ≈ 0ms. Surrogate spike trains preserving the bursting dynamics were used to remove this peak by subtracting the mean of the surrogate ISIHs, and the significance of the remaining peaks was assessed by the standard deviation of the surrogate ISIHs [63].

Surrogate spike trains were generated for each neuron by randomly swapping adjacent interspike intervals Δ*t_i_* = *t_i_* − *t*_*t*−1_ with 1 ≤ *i* ≤ *m*. The idea was to choose a random index 1 ≤ *i* ≤ *m*, swap the neighbouring inter-spike intervals Δ*t_i_* ↔ Δ*t*_*i*+1_, repeat this *ml* times, and add up the inter-spike intervals 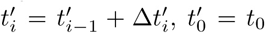 to obtain a surrogate spike train. The new series of inter-spike intervals was locally shuffled 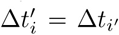 and the displacement *i*′ − *i* follows a binomial distribution with variance *σ*^2^ = *l*. Thus the temporal structure of the spike train was only locally disturbed, preserving both the inter spike interval histogram and a non-stationary spike rate (bursting).

For each pair of neurons, surrogate spike trains (*N* = 20, with *l* = 2) were created and the within-bin mean 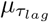 and standard deviation 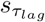 of their histograms were calculated. These were used to transform the original histogram *P* into a standard score *z* using the mean and standard deviation of the surrogate data, according to

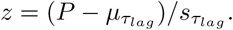

If the peak *z_max_* located at *τ_spike_* of this standard score, exceeded a fixed threshold, *z_max_* > *ρ* (Figure 5D, red peak above dashed line), we assumed a “functional connection” (instead of a “synapse”) with a defined timing between pre- and post-synaptic spike, *τ_spike_*.

Note, that the probability of observing a significance *z* or larger, if the null hypothesis is true, is given by 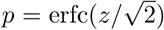 [63]. Thus *z* values for all pairwise connections can be transformed to *p* values.

**Figure 5:**
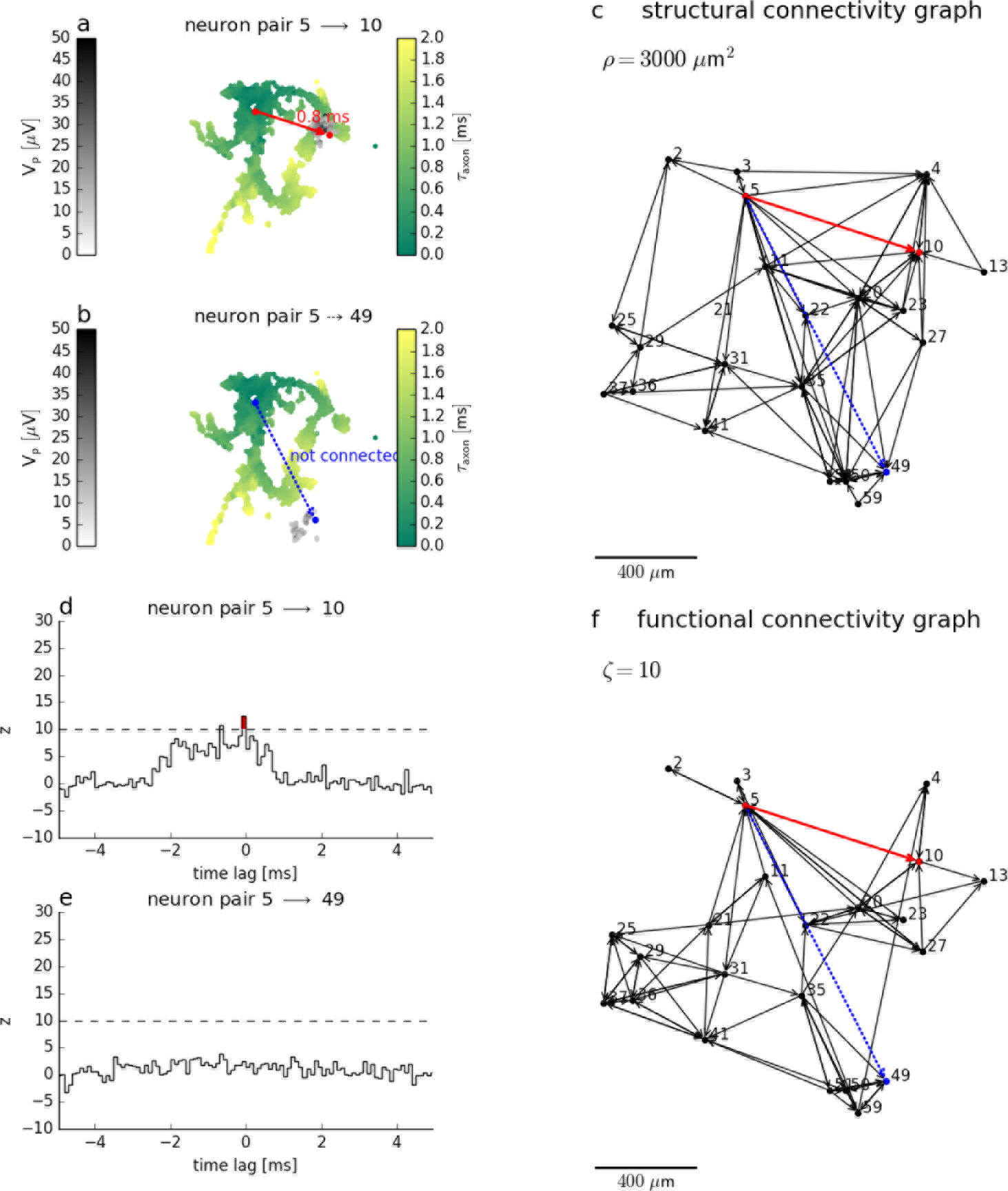
Estimation of structural and functional connectivity for a small network of *n* = 23 neurons from the spontaneous activity of the individual neurons. Structural connectivity graph **(c)** for an threshold *ρ* = 3000 *µm*^2^ for the pairwise overlap of axonal and dendritic fields. From a total of *k* = *n*(*n* − 1) = 506 possible pairwise connections, two examples are shown representing a pair of connected **(a)** and a pair of unconnected **(b)** neurons. Functional connectivity graph **(f)** for an threshold *ζ* = 10 for the z-score obtained from inter-spike interval histograms. From a total of *k* = *n*(*n* − 1) = 506 possible pairwise connections, two examples are shown: for a pair of connected **(d)** and a pair of unconnected **(e)** neurons. Note, that structural and functional connectivity are estimated from the spatial extension of spike-triggered averages and the temporal interdependence of individual spike trains, respectively. Although the examples show a match in structural and functional connectivity, the network graphs differ in a significant number of connections. Network layout **(c, f)** is plotted with respect to the position of the axon initial segments of its neurons.

### 2.5 Comparison of structural and functional connectivity

First, the pairwise, directed, structural and functional connectivity between all recorded neurons was estimated. The structural and functional connectivity graphs *S* and *F* of n neurons were represented by two matrices of *n* × *n* elements with the element with index *ij* equal to 1, if the *i*-th neuron was connected to the *j*-th neuron and 0 otherwise. Like the pairwise connectivity, the estimated structural and functional connectivity of the network depends on the thresholds for overlap *ρ* and the threshold for the score *ζ*, respectively.

We calculated some characteristic parameters [20] of these graphs, including the average clustering-coefficients, *C_S_* and *C_F_*, the average shortest path lengths, *L_S_* and *L_F_*, and the average degrees, *D_S_* and *D_F_*.

The degree of a single neurons *d* is the number of its pre- and postsynaptic neurons. Therefore, the average structural and functional degree for all *n* neurons can be calculated by *D_S_* = *k_S_/n* and *D_F_* = *k_F_/n*, where *k_S_* and *k_F_* are the total number of connections, respectively.

The clustering coefficient of a single neuron *c* measures how much the pre- and post-synaptic neurons of neurons are connected themselves. It is calculated by counting the triangles *k* that a neuron form with its connected neurons. This number is then divided by the maximum number of these triangles to obtain a value between 0 and 1:

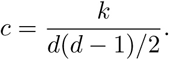

This value is 0, if pre- and post-synaptic neurons are not connected to each other, and it is 1 if they are all connected to each other. The average clustering coefficient *C* is obtained by averaging over the individual clustering coefficients *c* of all neurons in the network.

Looking at any two neurons in the network, in most cases the two neurons are not directly connected and there are several synapses between them. Here we use the estimated functional or structural connections as proxy for ‘synapses. The minimal number of the “synapses”, *l* that are necessary to link between the two neurons is also called the shortest path length. The characteristic path length, or average shortest path length, *L* is the average of the shortest path length *l* between any two connected neurons in the network. And in case the network consists of two or more, disconnected, smaller networks, we calculate *L_S_* and *L_F_* for the largest of these sub-networks.

Structural connectivity was calculated from the overlap of axonal and dendritic fields between all neurons. Although some axons extended over the whole recording area, most of them were rather short, representing a few long range and mostly short range connections. Therefore, some neurons could be structurally connected, whereas others could not be structurally connected, because they were well separated. An example connectivity graph for a rather high threshold on the overlap area of *ρ* = 3, 000 *µm*^2^ (corresponding to 10 electrodes) is shown in Figure 6c, with *k_S_* = 75. For *n* = 23 neurons the maximum number of unidirectional connections between two neurons, would amount to *k* = *n*(*n* − 1) = 506. Instead, we observed only 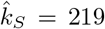 connections even for the lowest possible threshold (corresponding to 1 electrode). Therefore, the information provided by determining the axonal and dendritic arbours can exclude 57% of all possible connections. More connections can be excluded by raising the threshold for detection, as the number of structural connections *k_S_* decreases nearly linearly with log *ρ*. This is followed by a similar decrease in the average degree (Figure 6a). In contrast, the average minimum path length showed a peak 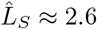 for an overlap threshold around *ρ* ≈ 1000 *µm*^2^. The average clustering *C_S_* ≈ 0.7 of each neuron was constant below that threshold.

**Figure 6.**
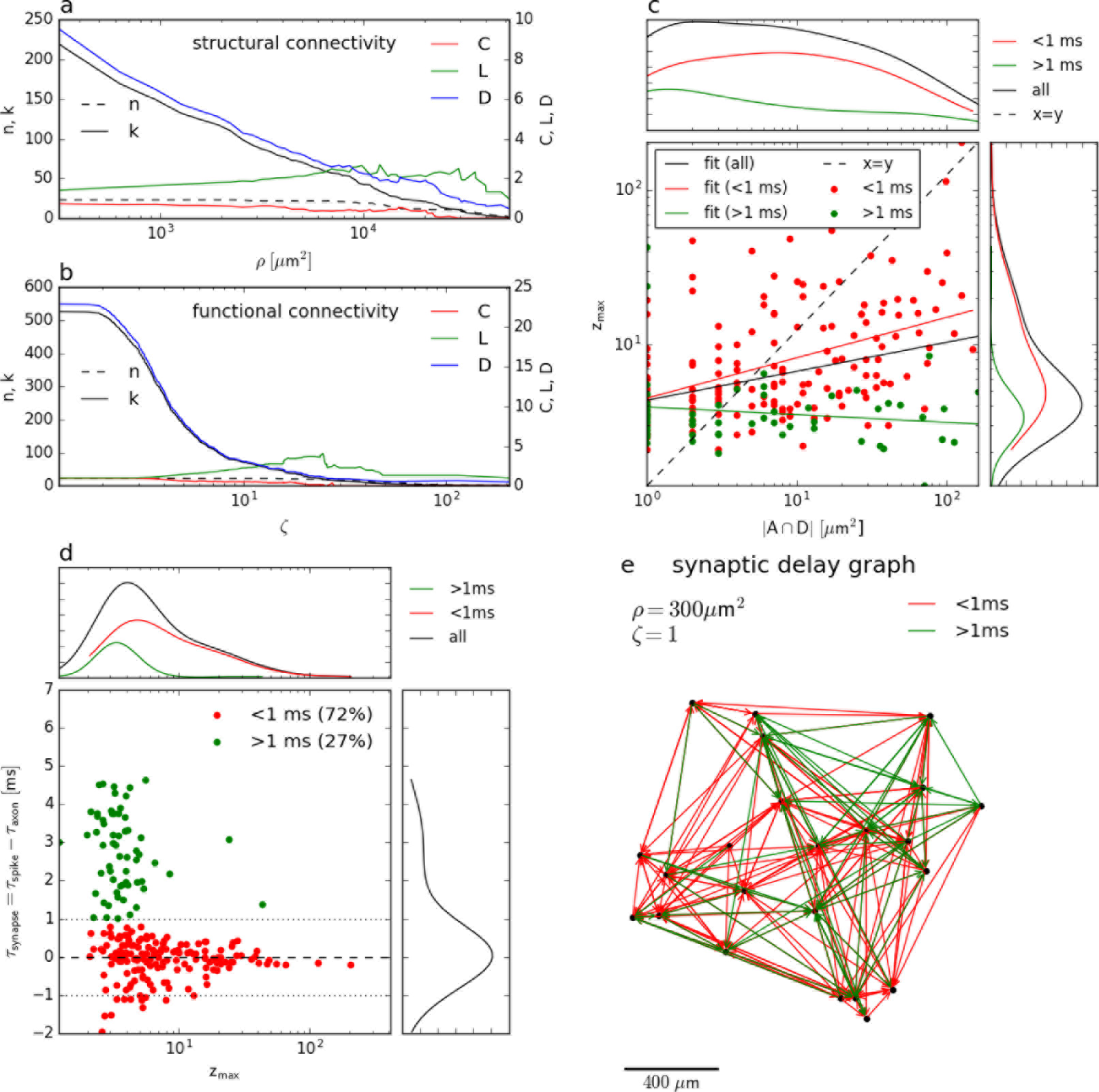
Comparison of structural and functional connectivity for a small network of *n* = 23 neurons. The number of edges (*k*), the clustering-coefficient (*C*), characteristic path length (*L*), and the average degree (*D*) of structural **(a)** and functional **(b)** connectivity graphs depend on the thresholds, *ρ* and *ζ*, respectively. Both graphs overlap, and this overlap is largest for the smallest thresholds (*ρ* = 300 *µm*^2^ and *ζ* = 2). Each connection is described by the strength (overlap between axon and dendrites, *A* ∩ *D*, vs. z-score, *z_max_*) and the delay (axonal delay, *τ_axon_*, vs. spike timing, *tau_spike_*). Surprisingly, the strength of structural and functional connections is only weakly correlated **(c)** with Pearson’s correlation test on log-transformed values, n=219, p<0.001, r=0.27 for all (black), n=66, p=0.02, r=−0.28 for delayed (green), and n=153, p<0.001, r: 0.41 for simultaneous (red) connections). Scatterplot **(d)** and graph **(e)** for synaptic delays (*τ_synapse_*) and z-scores (*z_max_*) reveal a network (green) of neurons connected by presumptive chemical synapses (*τ_synapse_* > 1 ms) with lower than average functional strength (median, 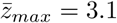). However for most connection the spike occurs almost instantaneous (*τ_axon_* = *τ_spike_*) and with greater reliability (median, 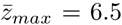). The difference in strength *z* is significant (Mood’s median test, *χ*^2^ =58.3, p<0.001, n=149 vs. 70).

Functional connectivity was accessed using a binning method and by that definition each neuron pair has such a prominent delay *τ_spike_*. However, the corresponding maximum scores have a rather large range, 2 ≤ *z_max_* ≤ 110. Thus for a low threshold of *ζ* < 2, all neurons were functionally connected, hence *k_F_* = 506, but for *ζ* > 3 the estimate of functional connectivity decreased (Figure 6b). This was accompanied by a increase in average minimum path length, which showed a clear peak at 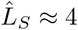 for a z-score threshold of *ζ* ≈ 20.

### 2.6 Estimation of synaptic delays

The first attempt to compare the structural and functional connectivity is to compare the strength of corresponding connections, whether a large overlap in axonal and dendritic fields is correlated with a stronger functional connectivity. Surprisingly, this correlation is weak (Figure 6c). The second attempt is to compare the delays, thus comparing the axonal delays with the spike timings (Figure 6d). Assuming chemical synapses, the prominent delay between two spike trains is a result of both, axonal conduction and the synaptic delay, caused by neurotransmitter release, diffusion, binding and action, thus *τ_spike_* = *τ_axon_* + *τ_synapse_*. This can be used to calculate the synaptic delay of presumably chemical synaptic connections,

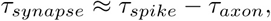

because we could measured both, the spike timing and the axonal delay. Most of the presumably synaptic delays were clustered around *τ_synapse_* ≈ 0 ms (red dots, Figure 6d), which means that these were either coincident spikes or caused by electric synapses which do not rely on neurotransmitters. A substantial number of delays were rather small, *τ_synapse_* ≪ 1 ms. Additionally, their functional score, *z_max_* is lower than that of “zero delay” functional connections, which is consistent with the lower reliability of chemical synapses.

### 2.7 Search for polychronous groups

The chemical synapses of cortical neurons are known to exhibit spike-timing-dependent plasticity (STDP) at millisecond precision[38]. The connecting axons conduct action potentials at millisecond precision[60]. Therefore, it has been hypothesised that these networks exhibit reproducible time-locked, but not synchronous firing patterns with millisecond precision, which are called polychronous groups[27].

To find polychronous groups, the possible (structural and functional) connections were restricted to those with putative chemical synapses. All spikes that occurred within the admissible delay

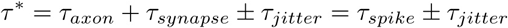

were connected and many connected spikes formed a spike pattern of a putative polychronous group. The spike pattern size varies from 2 to over 1000 spikes, and different neurons participated in the generation of these spike patterns.

In order to see how many spike patterns could be expected by chance this procedure was repeated for surrogate time series (see functional connectivity) as well as surrogate networks (Figure 7a). In the first surrogate network, the post-synaptic neurons were shuffled, but the number of post-synaptic neurons for each neuron and the delays of these connections *τ*\* (which are mainly determined by its axons) were preserved. In the second surrogate network, all connections were preserved, but the delays *τ*\* were shuffled within the network. All surrogates showed the same size distribution for the spike patterns, for a total of 7 cultures examined (see supplement), so that it could be concluded that the detected patterns arose just by chance (Figure 7d).

**Figure 7:**
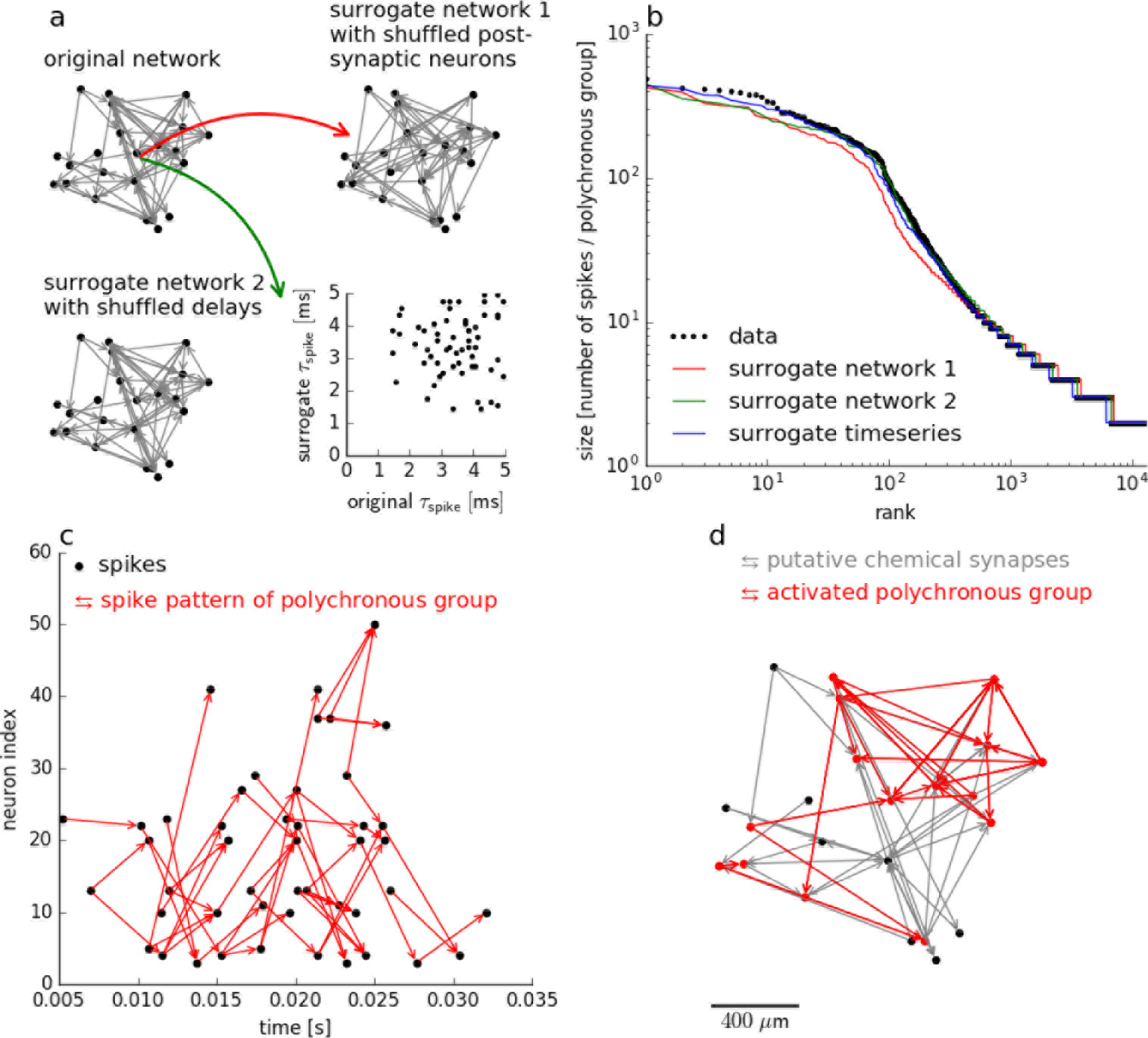
Absence of polychronisation in cultured neocortical neurons. Spike patterns consistent with measured axonal delays as well as synaptic delays of chemical synapses between the individual neurons were detected. Two different methods were used to generate surrogate networks **(a)**, together with surrogate spike trains. All surrogates showed the same size distribution for the putative polychronous groups **(b)**, thus the spike patterns detected in the original data arise just by chance. An example for a putative polychronous group is shown as scatterplot **(c)** of connected spikes. Only a part (red) of the putative chemical synapses (grey) participated in this spike pattern**(d)**.

### 2.8 Distinguishing neuron subtypes

Some classes of inhibitory neurons can be distinguished from excitatory neurons according to their spike waveform. For the electrode closest to the proximal fraction of the AIS the extracellular spikes always contained a large narrow negative spike, presumably caused by Na+ currents, followed by a wider positive peak, at least partially caused by potassium currents during the repolarization phase of the action potential [56][19]. This can be quantified by the delay between the negative peak and the positive peak *δ_p_*, also termed peak-to-peak width [14]:

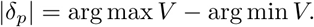

For fast spiking GABAergic neurons the latter is very short, |*δ_p_* | < 0.350*ms* [52]. All recordings were performed >28 DIV, when the chloride ion transporter KCC2 was already highly expressed in glutamatergic neurons, which decreased their intracellular chloride ion concentration [58], and therefore GABAergic neurons generally act as inhibitory neurons [51]. In our cultures, all recorded neurons showed a peak width |*δ_p_* | > 0.350*ms* so that they presumably represented excitatory, glutamatergic neurons. This was consistent with results obtained by GABA immunolabelling showing that the amount of GABAergic neurons was very low (<5%) (data not shown).

## 3 Discussion

We have presented a general framework to infer structural and functional connectivity to study emergent network behaviour. Although recording from axons and dendrites requires close proximity to the recording electrodes and therefore our methods is limited to neurons cultured on top of HDMEAs, it offers a novel and unique approach to study neuronal networks.

### 3.1 “Electrical imaging” of neuronal morphology

HDMEAs with more than 3,000 electrodes per square millimetre and dedicated low-noise on-chip amplifiers are a suitable tool to reveal the single-cell morphologies of cultured neurons.

We use HDMEAs to segment an electrical footprint into neuronal compartments (AIS, axonal arbours, dendritic arbour) or background activity according to the waveforms of the extracellular field potentials [47]. Typically, such information is not available or can only be obtained for a few cells by either intracellular dye injection [49], sparse transfection [4], or the Brainbow technique [36]. On the other hand, the dense reconstruction of neuronal networks from serial electron microscopic sections [7][32] is still very challenging.

We validated our method by comparing the axon segmentation with the ground truth morphology obtained by sparse transfection [4]. Our method works well for axons, because (a) they actively propagate action potentials and (b) they extend over large distance (≫ 200 *µ*m) without densely covering large areas of the array. This yields point-source-like signals that are clearly distinct from the background noise. In the case of dendrites only a rough outline can be obtained, because (a) they reveal themselves only by passive properties as the return current, (b) the extensive branching densely covers a small area, and (c) their signal is obscured by the action potential originating at the AIS, which is - at the same time - the very source of the return current. To some extent (b) could be resolved by increasing the electrode density.

Similar to optical imaging, there is a “resolution limit” for the electrode density of HDMEAs. The point spread function (PSF) for electrical recordings [35], can be calculated by

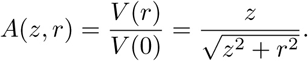

where *z* is the vertical distance of the signal source from the HDMEA and *r* the radial distance from the foot point. By solving *A*(*z*, *d_h_*/2) = 1/2 we find that the half-width 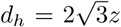 of this PSF depends on the distance *z*. This distance is very small for planar cell cultures (*z* = 3*µ*m) so that the maximum resolution is about *d_h_* = 10 *µ*m, which is about half the electrode distance in the HDMEA that was used in this study.

An optimised recording scheme for the switch matrix HDMEA design uses combinations of fixed and variable recording sites for high-throughput parallel mapping of the entire network of neurons. The method described here shows a promising way to obtain axonal arbours at large scale and potentially from all neurons during a single recording session.

### 3.2 Estimating both structural and functional connectivity

HDMEAs can be used to investigate how structural and functional connectivity are interrelated: Is functional connectivity present in the absence of structural connectivity? Is structural connectivity predictive for functional connectivity? The difficulty in answering these questions is the difficulty to obtain both, structural and functional connectivity for the very same experimental subject. There have been considerable achievements on the macroscopic level (whole brain) using a combination of fibre tracking through diffusion tensor imaging with mapping of macroscopic (brain metabolic) activity by BOLD imaging [57][25]. On the mesoscopic level, the use of various all-optical methods has been put forward, but the combination of optical tracers with imaging of activity-dependent calcium signals and optical stimulation is difficult due the limited number of discernible wavelengths and limited temporal resolution.

Here we propose an all-electrical method to estimate the structural connectivity according to a common extension of Peter’s rule [46][8], which assumes that the probability of two neurons being connected is proportional to the overlap of their axonal and dendritic fields [50]. Currently, this rule is controversial and refuting [55][41][10][29][32] and confirming [48][22] evidence has been reported.

At the same time we used pairwise inter-spike interval histograms for the estimation of the functional connectivity. The significance has been accessed by surrogate time series. This approach is more general than previous methods, using bin-less methods [40] or including higher order interactions [6].

### 3.3 Testing a spike-based theory of neuronal computation

Theories of neuronal computation are usally divided into two types [9]: “Rate-based” theories are based on the firing rate, for example rank order coding [64] and predictive spike coding [15], whereas “spike-based” theories propose some kind of synchrony between neurons leading to repeated spike patterns, such as synfire chains [1] and polychronization [27]. Polychronisation refers to the theoretical phenomenon that recurrent networks with axonal delays and spike timing depended plasticity (STDP) [38] exhibit complex activity patterns [27]. These precisely timed spikes are produced by so called polychronous groups of neurons for which the combination of axonal delays is just right to favour their sustained recurrent activation. In simulations, these groups form both spontaneously and in response to patterned stimulation, which has been used as an argument that this phenomenon might underlie memory processes [61].

For the first time, the direct measurement of axonal delays between each neuron in the network and the identification of chemical synapses allowed us to experiment on this popular spike-based theory of neuronal computation. Here we can show that spike patterns consistent with axonal delays between pre- and post-synaptic neurons arise just by chance, thus favouring rate-based neuronal computation.

Another explanation lies in the difficulty to observe STDP in bursting cultured neuronal networks. Whereas numerous studies demonstrate that neuronal networks show short-term plasticity (elasticity) upon single stimulations in cultures [16], the reports regarding long term effects have been conflicting [67]. Although we found that some synaptic delays are consistent with chemical synapses observed in dissociated cortical neuron cultures [43], the exact nature and plasticity of a synaptic connection can only be examied by combining single-cell stimulation of the pre-synaptic cell with intracellular recording of post-synaptic potentials [28].

### 3.4 High-throughput electrophysiology of full axonal arbours

The presented method to automatically segment axons of a large number of individual neurons also holds potential for high-throughput single cell applications to study the physiology of full axonal arbours [2][53][30][31][13]. Up to now, axonal physiology has been investigated using either very large axons (giant squid axon) [24][26] or paired patch recordings that are possible if either naturally or experimentally induced swelling enables patching of an individual axonal branch [54].

Axonal signals have been recorded extracellularly with standard planer multi-electrode arrays (MEAs) [3][65], but their spatial resolution is limited by the larger distance of the electrodes in MEAs (*r* = 100*µ*m) compared to HDMEAs (*r* = 18*µ*m), which allow the reconstruction of axonal arbors [4]. It has been argued, that HDMEAs produce very large data files that may limit the high-throughput use of this technology as well as limit the increased signal to noise obtained by averaging many hundreds of waveforms obtainable in a longer recording [65]. Here we show that we can isolate full axonal arbors of up to 52 neurons in parallel from a single two hour recording session (see Supplemental figure 1 for more examples). This time can be further improved using a HDMEA design with increased number of recording channels [42], decreasing the average number of configurations per neuron to *C* = 4*e/a*^2^ = 4 × 26400/10242 = 0.1 (see section 2.1). With a recording time of 2 min/configuration for spike-triggered averaging the full axonal arbors of 500 neurons can be reconstructed from a single 2 hour recording session.

Although our method is limited to neurons in planar culture, it offers the advantage of recording (a) at the entire axonal arbour from the AIS to the terminal branches and everywhere in between and (b) from several neurons in parallel. The criss-cross pattern of axonal branches might even be advantageous when combined with local application of drugs, because even for local application several branches of different neurons can be screened in parallel. A motorised micro-manipulator [35] can be used map out the distribution of a specific ion channel [44] over many individual neurons by using a specific inhibitor in a single experiment.

### 3.5 Conclusion

Our all-electrical framework for neuron cultures on a planar HDMEAs enables the estimation of structural connectivity along with an investigation of functional coupling. Our data show that the “structural” connectivity for a small network overlaps with the “functional” connectivity obtained from the pairwise inter-spike interval histograms. For 31% of these connections, the estimated synaptic delay was consistent with that of chemical synapses [43]. We did not find any evidence for polychronisation [27].

Currently, our cortical preparation is quite artificial, as it contains most excitatory neurons, but using preparations of other brain areas (e.g., striatum) any excitatory-inhibitory balance (E/I) can be obtained and its functional significance can be investigated [12]. Furthermore, the lack of input from thalamic areas [33] drives the activity into the bursting regime reminiscent of sleep. Chemical stimulation [23] or patterned focal stimulation can be used to induce an active wake-like state exhibiting random sparse firing. This would make it possible to study more natural properties of cortical networks closer to the cortex of behaving animals [11].

Interestingly, neuron dynamics can be reduced in most cases to a leaky integrate-and-fire (LIF) model with only a few free parameters, which can, together with measured axonal delays, be used to reconstruct the network *in silico* [68]. Comparing the activity of these simulated networks with actual activities of networks from which they were reconstructed, would provide a “test-bed” for the validation of recent large-scale cortical simulations.

In general, our method can be used to investigate the relationship between the topology of neuronal connections and emerging temporal spike patterns observed in dissociated neuronal cultures. This new framework can be used to test a wide range of rate-based vs. spike-based theories of neuronal computation as well as to evaluate axonal and synaptic plasticity in neuronal networks.

## 4 Methods

### 4.1 Animal use

Timed pregnant rats (Wistar) were obtained from a commercial vendor (Nihon SLC, Japan). Animals were sacrificed on the day of arrival to obtain embryos for primary neuron cultures. All experimental procedures on animals were carried out in accordance with the European Council Directive of 22 September 2010 (2010/63/EU) and had been approved by the local authorities (Animal Care and Use Committee of RIKEN; QAH24-01).

### 4.2 High-density microelectrode array (HDMEA)

Sub-cellular resolution extracellular recordings were obtained using a complementary metal-oxide-semiconductor (CMOS)-based HDMEA [18] with 11,011 electrodes arranged in a hexagonal pattern yielding an electrode density of 3,150 electrodes/mm^72^. The culture chamber of the HDMEA was prepared as described before [21] with minor modifications: after attaching the chamber ring (polycarbonate, 19 mm inner diameter, 8 mm high) using epoxy resin (EPO-TEK 301-2, Epoxy Technology Inc.), GlobTop was used to cover the bond wires while keeping the electrode area clean, and the remaining area was covered by a thin film of PDMS (Sylgard 184, Dow Corning). Platinum black was electrochemically deposited [39] with modification by [21] on the electrodes to decrease their impedance in order to reduce recording noise and increase signal gain [66]. One day before plating, the surface of the HDMEAs was rendered hydrophilic by oxygen plasma treatment (40s, 20W), coated for 4 hours with of 50 *µ*g/ml Poly-D-Lysine (Sigma-Aldrich, P7280) in PBS, washed twice with Aqua dest. and air-dried for one hour.

### 4.3 Primary neuron cultures

Adult rats were anaesthetised with Isofluorane and killed using a guillotine. The embryos were removed from the uterus and decapitated. Their neocortex was dissected in ice-cold dissection medium (HBSS without Ca2+ and Mg2+; Gibco, NO. 14175) and incubated for 20 minutes at 37*°C* in Trypsin/EDTA (Sigma-Aldrich). After washing twice with plating medium (Neurobasal A supplemented with 10% Fetal bovine serum, 2% B27 Supplement, 1:100 GlutaMax, all from Gibco, and 10 *µ*g/ml Gentamicin, Sigma-Aldrich), the tissue was mechanically dissociated, passed through a 40 *µ*m nylon mesh, and centrifuged 6 min at 200 g. The supernatant was removed; the cells were suspended and counted. A 20 *µ*l drop containing 10,000 cells was placed on the electrode area of the HDMEA in the middle of the culture chamber. The cultures were covered by a membrane permeable to gas but not to water vapour (Potter and DeMarse, 2001) and placed in a standard incubator (37°*C*, 5% CO2, 80% relative humidity). The neurons were allowed to settle and attach to the surface for 30 minutes. Thereafter, the culture chamber was filled with 600 *µ*l serum-free, astrocyte conditioned DMEM/Hams’s F12 medium (Nerve Culture Medium, Sumitomo, #MB-X9501). Medium was exchanged completely with 600 *µ*l conditioned medium after 4 days and then every 7 days.

### 4.4 Statistical significance

A total of 8 cultures from 3 different neuron preparations were examined. For 7 cultures the structural and functional connectivity was estimated; the networks consisted of 14–52 neurons/culture (see supplemental figure 1). One culture was transfected with an expression plasmid for DsRed, and the spike-triggered averages for a single, isolated, DsRed-expressing neuron were extracted from the recordings.

### 4.5 Live imaging

Live-cell visualisation of whole neurons was performed by transfection [4] [49]. Transfection was performed using a pLV-hSyn-RFP plasmid from Edward Callaway (Addgene, #22909) and Lipofectamine 2000 (Life Technologies) in accordance with the manufacturer’s protocol. A Leica DM6000 FS microscope, Leica DFC 345 FX camera, and the Leica Application Suite software were used to produce micrographs.

### 4.6 Recordings

For recording, HDMEAs were place in a bench-top incubator (TOKAI HIT, INU-OTOR-RE) with temperature control, and 5% CO2 was supplied by a gas-mixer and humidified by a water bath. In order to avoid the evaporation of medium during prolonged recording intervals, the water bath and the lid temperature set point was set 1°*K* and 3°*K* above the sample temperature set point, which was 35°*C*.

The HDMEA was attached to the adapter, and the recordings were performed using custom scripts written in LabView (National Instruments), Matlab (Mathworks), C++ and Python running on a standard PC with a Linux operating system. Data underwent lossless data compression and were directly stored on a server on the local LAN.

Offline analysis of the recordings included filtering, event detection and averaging. First, a band-pass filter (2nd order Butterworth filter, 100-3500 Hz) removed slowly changing field potentials as well as high frequency noise. The remaining (background) noise was characterised by the median absolute deviation (MAD), which is resilient to outliers in the data but represents a consistent estimator of the standard deviation, *s_V_* = 1.4826 MAD(*V_sig_*). Using a voltage threshold method for event detection [34], negative peaks below a threshold of *V_thr_* = 6 × *s_V_* (thus *V_thr_* ≫ 50*µV*) were identified. To avoid multiple detection of the same spike, successive events within less than 0.5 ms were discarded. To extract sub-threshold events from axons and dendrites, signal averaging was performed (see results).

## 5 Additional information

### 5.2 Author Contributions

TB planned experiments, performed cell culture, recordings, implemented and tested algorithms in Matlab and Python, performed data analysis, prepared figures, discussed results and wrote manuscript; MR performed cell culture, recordings, transfection, imaging and discussed results; SH implemented and tested algorithms for data analysis in Python, discussed results; KD prepared neocortical neurons, performed immunostaining, imaging, discussed results; AH provided HDMEAs, discussed results; UF planned experiments, discussed results, implemented software for recording and data analysis and wrote manuscript.

### 5.3 Supplementary Material

The Hana (<u>h</u>igh density microelectrode array recording <u>ana</u>lysis) analysis pipeline is open source. All source code as well as example data to replicate the figures will be available at Github. The example data consists of spike triggered-averages and events that were extracted from the raw recordings.

## Supplemental figures legends

Supplemental figure 1. Summaries for the analysis of connectivity in spontaneous active neuronal networks (n=7) cultured on HDMEA. Structural connectivity graph **(a)** for an threshold *ρ* = 300 *µm*^2^ for the pairwise overlap of axonal and dendritic fields. Functional connectivity graph **(b)** for an threshold *ζ* = 1 for the z-score obtained from inter-spike interval histograms. Synaptic delay graph **(c)** revealing a network (green) of neurons connected by presumptive chemical synapses (*τ_synapse_* > 1 ms) and another for which which the spike occurs almost instantaneous (*τ_axon_* = *τ_spike_*). Correlation plots for strength of structural and functional connections **(d)** for all (black), for delayed (green), and for simultaneous (red) connections. Scatterplot **(e)** for synaptic delays (*τ_synapse_*) and z-scores (*z_max_*) reveal a network (green) of neurons connected by presumptive chemical synapses (*τ_synapse_* > 1 ms) and other connection for which the spike occurs almost instantaneous (*τ_axon_* = *τ_spike_*). Size distribution for the spike patterns detected in the original data **(f)** together is the same as for the surrogates. Thus the original spike patterns similarly arise just by chance and not from the polychronous groups. Note: The networks **(a, b, c)** are plotted with respect to the position of the axon initial segments of its neurons.

Supplemental figure 2. Axonal (blue) and dendritic (red) fields of all spontaneous active neurons in a small neuronal network (culture 1) reconstructed from the HDMEA recording.

## References

1. M. Abeles. Corticonics: Neuronal Circuits of the Cerebral Cortex. Cambridge University Press, Cambridge, England, 1st edition, 1991.

2. Henrik Alle and Jorg R P Geiger. Combined analog and action potential coding in hippocampal mossy fibers. Science, 311(5765):1290–1293, Mar 2006.

3. Douglas J Bakkum, Zenas C Chao, and Steve M Potter. Long-term activity-dependent plasticity of action potential propagation delay and amplitude in cortical networks. PLOS one, 3(5):e2088, 2008.

4. Douglas J Bakkum, Urs Frey, Milos Radivojevic, Thomas L Russell, Jan Müller, Michele Fiscella, Hirokazu Takahashi, and Andreas Hierlemann. Tracking axonal action potential propagation on a high-density microelectrode array across hundreds of sites. Nat Commun, 4:2181, Jan 2013.

5. Douglas J. Bakkum, Milos Radivojevic, David Jaeckel, Felix Franke, Thomas L Russell, Urs Frey, Hirokazu Takahashi, and Andreas Hierlemann. The axon initial segment is a strong contributor to a neuron’s local extracellular field potential. In Society for Neuroscience (SfN) Conference, number 1, page 372.05, Washington DC, 2014.

6. Luís M A Bettencourt, Greg J Stephens, Michael I Ham, and Guenter W Gross. Functional structure of cortical neuronal networks grown in vitro. Phys Rev E Stat Nonlin Soft Matter Phys, 75(2 Pt 1):021915, Feb 2007.

7. Davi D Bock, Wei-Chung Allen Lee, Aaron M Kerlin, Mark L Andermann, Greg Hood, Arthur W Wetzel, Sergey Yurgenson, Edward R Soucy, Hyon Suk Kim, and R Clay Reid. Network anatomy and in vivo physiology of visual cortical neurons. Nature, 471(7337):177–182, 2011.

8. Valentino Braitenberg and Almut Schütz. Cortex: Statistics and geometry of neuronal connectivity. Buch, pages 1–118, Sep 2010.

9. Romain Brette and Emmanuel Guigon. Reliability of spike timing is a general property of spiking model neurons. Neural Computation, 15(2):279–308, 2003.

10. Kevin L Briggman, Moritz Helmstaedter, and Winfried Denk. Wiring specificity in the direction-selectivity circuit of the retina. Nature, 471(7337):183–188, 2011.

11. Zenas C Chao, Douglas J Bakkum, Daniel A Wagenaar, and Steve M Potter. Effects of random external background stimulation on network synaptic stability after tetanization: a modeling study. Neuroinformatics, 3(3):263–80, 2005.

12. Xin Chen and Rhonda Dzakpasu. Observed network dynamics from altering the balance between excitatory and inhibitory neurons in cultured networks. Physical Review E, 82(3):031907, 2010.

13. Dominique Debanne. Information processing in the axon. Nat Rev Neurosci, 5(4):304–16, Apr 2004.

14. Kosmas Deligkaris, Torsten Bullmann, and Urs Frey. Extracellularly recorded somatic and neuritic signal shapes and classification algorithms for high-density microelectrode array electrophysiology. Frontiers in Neuroscience, 10:421, 2016.

15. Sophie Deneve. Bayesian spiking neurons i: inference. Neural computation, 20(1):91–117, 2008.

16. Danny Eytan, Naama Brenner, and Shimon Marom. Selective adaptation in networks of cortical neurons. Journal of Neuroscience, 23(28):9349–9356, 2003.

17. Tom Fawcett. An introduction to roc analysis. Pattern recognition letters, 27(8):861–874, 2006.

18. Urs Frey, Jan Sedivy, Flavio Heer, Rene Pedron, Marco Ballini, Jan Mueller, Douglas Bakkum, Sadik Hafizovic, Francesca D Faraci, Frauke Greve, et al. Switch-matrix-based high-density microelectrode array in cmos technology. Solid-State Circuits, IEEE Journal of, 45(2):467–482, 2010.

19. Carl Gold, Darrell A Henze, Christof Koch, and György Buzsáki. On the origin of the extra-cellular action potential waveform: A modeling study. J Neurophysiol, 95(5):3113–28, May 2006.

20. Suchin S Gururangan, Alexander J Sadovsky, and Jason N MacLean. Analysis of graph invariants in functional neocortical circuitry reveals generalized features common to three areas of sensory cortex. PLoS Comput Biol, 10(7):e1003710, Jul 2014.

21. F. Heer, S. Hafizovic, W. Franks, A. Blau, C. Ziegler, and A. Hierlemann. Cmos microelectrode array for bidirectional interaction with neuronal networks. IEEE Journal of Solid-State Circuits, 41(7):1620–1629, July 2006.

22. Sean L Hill, Yun Wang, Imad Riachi, Felix Schürmann, and Henry Markram. Statistical connectivity provides a sufficient foundation for specific functional connectivity in neocortical neural microcircuits. Proc Natl Acad Sci U S A, 109(42):E2885–94, Oct 2012.

23. Valérie Hinard, Cyril Mikhail, Sylvain Pradervand, Thomas Curie, Riekelt H Houtkooper, Johan Auwerx, Paul Franken, and Mehdi Tafti. Key electrophysiological, molecular, and metabolic signatures of sleep and wakefulness revealed in primary cortical cultures. J Neurosci, 32(36):12506–17, Sep 2012.

24. A L Hodkin and A F Huxley. A quantitative description of membrane current and its application to conduction and excitation in nerve. J Physiol, 117(4):500–544, Aug 1952.

25. C J Honey, O Sporns, L Cammoun, X Gigandet, J P Thiran, R Meuli, and P Hagmann. Predicting human resting-state functional connectivity from structural connectivity. Proc *Natl Acad Sci U S A*, 106(6):2035–40, Feb 2009.

26. A F Huxley. Ion movements during nerve activity. Ann N Y Acad Sci, 81:221–246, 1959.

27. Eugene M Izhikevich. Polychronization: computation with spikes. Neural Comput, 18(2):245–82, Feb 2006.

28. David Jäckel, Jan Müller, Thomas L. Russell Milos Radivojevic, Felix Franke, Urs Frey, Douglas J. Bakkum, and Andreas Hierlemann. Simultaneous Intra- and Extracellular Recordings using a Combined High-Density Microelectrode Array and Patch-Clamp System. In Proceedings MEA Meeting 2014: July 1 - July 4, 2014, Reutlingen, Germany: 9th International Meeting on Substrate-Integrated Microelectrode Arrays, volume 9, pages 153–155, Reutlingen, 2014. NMI Natural and Medical Sciences Institute at the University of Tuebingen.

29. Narayanan Kasthuri, Kenneth Jeffrey Hayworth, Daniel Raimund Berger, Richard Lee Schalek, Jose Angel Conchello, Seymour Knowles-Barley, Dongil Lee, Amelio Vazquez-Reina, Verena Kaynig, Thouis Raymond Jones, Mike Roberts, Josh Lyskowski Morgan, Juan Carlos Tapia, H Sebastian Seung, William Gray Roncal, Joshua Tzvi Vogelstein, Randal Burns, Daniel Lewis Sussman, Carey Eldin Priebe, Hanspeter Pfister, and Jeff William Lichtman. Saturated reconstruction of a volume of neocortex. Cell, 162(3):648–661, Jul 2015.

30. Maarten H P Kole. First node of ranvier facilitates high-frequency burst encoding. Neuron, 71(4):671–82, Aug 2011.

31. Maarten H P Kole and Greg J Stuart. Signal processing in the axon initial segment. Neuron, 73(2):235–47, Jan 2012.

32. Wei-Chung Allen Lee, Vincent Bonin, Michael Reed, Brett J Graham, Greg Hood, Katie Glattfelder, and R Clay Reid. Anatomy and function of an excitatory network in the visual cortex. Nature, 532(7599):370–374, 2016.

33. Maxime Lemieux, Jen-Yung Chen, Peter Lonjers, Maxim Bazhenov, and Igor Timofeev. The impact of cortical deafferentation on the neocortical slow oscillation. J Neurosci, 34(16):5689–703, Apr 2014.

34. M S Lewicki. A review of methods for spike sorting: the detection and classification of neural action potentials. Network, 9(4):R53–78, Nov 1998.

35. Jing Lin, Marie Engelene J Obien, Andreas Hierlemann, and Urs Frey. Automated navigation of a glass micropipette on a high-density microelectrode array. In Engineering in Medicine and Biology Society (EMBC), 2015 37th Annual International Conference of the IEEE, pages 881–884. IEEE, 2015.

36. Jean Livet, Tamily A Weissman, Hyuno Kang, Ryan W Draft, Lu Ju, Robyn A Bennis, Joshua R Sanes, and Jeff W Lichtman. Transgenic strategies for combinatorial expression of fluorescent proteins in the nervous system. Nature, 450(7166):56–62, Nov 2007.

37. H Lohmann and B Rörig. Long-range horizontal connections between supragranular pyramidal cells in the extrastriate visual cortex of the rat. Journal of Comparative Neurology, 344(4):543–558, 1994.

38. H Markram, J Lübke, M Frotscher, and B Sakmann. Regulation of synaptic efficacy by coincidence of postsynaptic aps and epsps. Science, 275(5297):213–5, Jan 1997.

39. Carl A Marrese. Preparation of strongly adherent platinum black coatings. Analytical chemistry, 59(1):217–218, 1987.

40. Mohammad Shahed Masud and Roman Borisyuk. Statistical technique for analysing functional connectivity of multiple spike trains. Journal of Neuroscience Methods, 196(1):201–219, Mar 2011.

41. Yuriy Mishchenko, Hu Tao, Josef Spacek, John Mendenhall, M Kristen. Harris, and B Dmitri. Chklovskii. Ultrastructural analysis of hippocampal neuropil from the connectomics perspective. Neuron, 67(6):1009–1020, Sep 2010.

42. Jan Müller, Marco Ballini, Paolo Livi, Yihui Chen, Milos Radivojevic, Amir Shadmani, Vijay Viswam, Ian L Jones, Michele Fiscella, Roland Diggelmann, Alexander Stettler, Urs Frey, Douglas J Bakkum, and Andreas Hierlemann. High-resolution cmos mea platform to study neurons at subcellular, cellular, and network levels. Lab Chip, May 2015.

43. Keiko Nakanishi and Fumio Kukita. Functional synapses in synchronized bursting of neocortical neurons in culture. Brain research, 795(1):137–146, 1998.

44. Zoltan Nusser. Variability in the subcellular distribution of ion channels increases neuronal diversity. Trends in Neurosciences, 32(5):267–274, May 2009.

45. Marie Engelene J Obien, Kosmas Deligkaris, Torsten Bullmann, Douglas J Bakkum, and Urs Frey. Revealing neuronal function through microelectrode array recordings. Frontiers in neuroscience, 8:423, 2015.

46. A Peters and M L Feldman. The projection of the lateral geniculate nucleus to area 17 of the rat cerebral cortex. i. general description. J Neurocytol, 5(1):63–84, Feb 1976.

47. Anders V Petersen, Emil Ø Johansen, and Jean-François Perrier. Fast and reliable identification of axons, axon initial segments and dendrites with local field potential recording. Front Cell Neurosci, 9:429, 2015.

48. T. C. Potjans and M. Diesmann. The cell-type specific cortical microcircuit: Relating structure and activity in a full-scale spiking network model. Cerebral Cortex, 24(3):785–806, Dec 2012.

49. Milos Radivojevic, David Jäckel, Michael Altermatt, Jan Müller, Vijay Viswam, Andreas Hierlemann, and Douglas J Bakkum. Electrical identification and selective microstimulation of neuronal compartments based on features of extracellular action potentials. Sci Rep, 6:31332, 2016.

50. Christopher L Rees, Keivan Moradi, and Giorgio A Ascoli. Weighing the evidence in peters’rule: Does neuronal morphology predict connectivity? Trends Neurosci, Dec 2016.

51. C Rivera, J Voipio, J A Payne, E Ruusuvuori, H Lahtinen, K Lamsa, U Pirvola, M Saarma, and K Kaila. The k+/cl-co-transporter kcc2 renders gaba hyperpolarizing during neuronal maturation. Nature, 397(6716):251–255, Jan 1999.

52. Ashlee A Robbins, Steven E Fox, Gregory L Holmes, Rodney C Scott, and Jeremy Michael Barry. Short duration waveforms recorded extracellularly from freely moving rats are representative of axonal activity. Frontiers in neural circuits, 7:181, 2013.

53. Takuya Sasaki, Norio Matsuki, and Yuji Ikegaya. Action-potential modulation during axonal conduction. Science, 331(6017):599–601, Feb 2011.

54. Takuya Sasaki, Norio Matsuki, and Yuji Ikegaya. Targeted axon-attached recording with fluorescent patch-clamp pipettes in brain slices. Nat Protoc, 7(6):1228–34, Jun 2012.

55. Gordon M G Shepherd, Armen Stepanyants, Ingrid Bureau, Dmitri Chklovskii, and Karel Svoboda. Geometric and functional organization of cortical circuits. Nat Neurosci, 8(6):782–90, Jun 2005.

56. Sergey Sitnikov, David Jaeckel, and Andreas Hierlemann. Studying extracellular action potential waveforms using hd meas. In Proceedings MEA Meeting 2014: June 28 - July 1, 2016, Reutlingen, Germany: 10th International Meeting on Substrate-Integrated Microelectrode Arrays, number 130, 2016.

57. Pawel Skudlarski, Kanchana Jagannathan, Vince D Calhoun, Michelle Hampson, Beata A Skudlarska, and Godfrey Pearlson. Measuring brain connectivity: diffusion tensor imaging validates resting state temporal correlations. Neuroimage, 43(3):554–61, Nov 2008.

58. Valentin Stein, Irm Hermans-Borgmeyer, Thomas J Jentsch, and Christian A Hübner. Expression of the kcl cotransporter kcc2 parallels neuronal maturation and the emergence of low intracellular chloride. Journal of Comparative Neurology, 468(1):57–64, 2004.

59. Olav Stetter, Demian Battaglia, Jordi Soriano, and Theo Geisel. Model-free reconstruction of excitatory neuronal connectivity from calcium imaging signals. PLoS Comput Biol, 8(8):e1002653, Jan 2012.

60. Harvey A Swadlow. Efferent neurons and suspected interneurons in motor cortex of the awake rabbit: properties axonal, sensory receptive fields, and subthreshold synaptic inputs. Journal of neurophysiology, 71(2):437–453, 1994.

61. Botond Szatmáry and Eugene M Izhikevich. Spike-timing theory of working memory. PLoS Comput Biol, 6(8), 2010.

62. Albert E Telfeian and Barry W Connors. Widely integrative properties of layer 5 pyramidal cells support a role for processing of extralaminar synaptic inputs in rat neocortex. Neuroscience letters, 343(2):121–124, 2003.

63. James Theiler, Stephen Eubank, André Longtin, Bryan Galdrikian, and J Doyne Farmer. Testing for nonlinearity in time series: the method of surrogate data. Physica D: Nonlinear Phenomena, 58(1-4):77–94, 1992.

64. Simon Thorpe, Arnaud Delorme, and Rufin Van Rullen. Spike-based strategies for rapid processing. Neural networks, 14(6):715–725, 2001.

65. Kenneth R Tovar, Daniel C Bridges, Wu Bian, Connor Randall, Morgane Audouard, Jiwon Jang, Paul K Hansma, and Kenneth S Kosik. Recording action potential propagation in single axons using multi-electrode arrays. bioRxiv, 2017.

66. Vijay Viswam, David Jäckel, Ian Jones, Marco Ballini, Jan Muller, Urs Frey, Felix Franke, and Andreas Hierlemann. Effects of sub-10*µ*m electrode sizes on extracellular recording of neuronal cells. In MicroTAS, 2014.

67. Daniel A Wagenaar, Jerome Pine, and Steve M Potter. Searching for plasticity in dissociated cortical cultures on multi-electrode arrays. J Negat Results Biomed, 5:16, 2006.

68. Yury V. Zaytsev, Abigail Morrison, and Moritz Deger. Reconstruction of recurrent synaptic connectivity of thousands of neurons from simulated spiking activity. Journal of Computational Neuroscience, 39(1):77–103, Jun 2015.

